# When is a bacterial “virulence factor” really virulent?

**DOI:** 10.1101/061317

**Authors:** Elisa T. Granato, Freya Harrison, Rolf Kümmerli, Adin Ross-Gillespie

**Affiliations:** Department of Plant and Microbial Biology, University of Zurich, Zurich, Switzerland; Centre for Biomolecular Sciences, University of Nottingham, Nottingham, United Kingdom

**Author notes:** Address correspondence to Elisa T. Granato. Present address: Adin Ross-Gillespie, SIB Swiss Institute of Bioinformatics, 13 Lausanne, Switzerland.

## Abstract

Bacterial traits that contribute to disease are termed ‘virulence factors’ and there is much interest in therapeutic approaches that disrupt such traits. However, ecological theory predicts disease severity to be multifactorial and context dependent, which might complicate our efforts to identify the most generally important virulence factors. Here, we use meta-analysis to quantify disease outcomes associated with one well-studied virulence factor – pyoverdine, an iron-scavenging compound secreted by the opportunistic pathogen *Pseudomonas aeruginosa*. Consistent with ecological theory, we found that the effect of pyoverdine, albeit frequently contributing to disease, varied considerably across infection models. In many cases its effect was relatively minor, suggesting that pyoverdine is rarely essential for infections. Our work demonstrates the utility of meta-analysis as a tool to quantify variation and overall effects of purported virulence factors across different infection models. This standardised approach will help us to evaluate promising targets for anti-virulence approaches.

## INTRODUCTION

Understanding which bacterial characteristics contribute most to disease is a major area of research in microbiology and infection biology (1–3). Bacterial characteristics that reduce host health and/or survival are considered ‘virulence factors’. Such factors include structural features like flagella and pili that facilitate attachment to host cells (4, 5), as well as secreted products like toxins and enzymes that degrade host tissue (6, 7), or siderophores that scavenge iron from the host (8). Research on virulence factors has not only increased our fundamental understanding of the mechanisms underlying virulence, but has also identified potential novel targets for antibacterial therapy. There is indeed much current interest in developing ‘antivirulence’ drugs to disrupt virulence factor production – the idea being that by simply disarming pathogens rather than killing them outright, we could ostensibly elicit weaker selection for drug resistance (9–11).

Although our understanding of different types of virulence factors and their interactions is continuously deepening, it is still unclear just how generalizable this assembled knowledge is. It is often assumed, for reasons of parsimony, that a given structure or secreted molecule central to the virulence of a particular bacterial strain in a specific host context will similarly enhance virulence in another bacterial strain, or in a different host (12). Yet, ecological theory predicts that the effects of a given trait will frequently vary in response to the environment (78). In the context of infections, this may be particularly true for opportunistic pathogens, which face very heterogeneous environments: they can live in environmental reservoirs (e.g. soil, household surfaces), as commensals of healthy hosts, or, when circumstances allow, as pathogens, causing serious infections in a range of different hosts and host tissues (13–15). Opportunistic pathogens underlie many hospital-acquired infections, especially in immune-compromised patients (16–18), and the treatment of such infections is often challenging because, as generalists, such pathogens are preselected to be tenacious and highly adaptable. Thus, in designing new anti-virulence drugs against opportunistic pathogens, we need to know not only whether the targeted trait is indeed associated with pathogenicity, but also the generality of this association across different pathogen strain backgrounds, host species, and infection types.

Here we show how a meta-analysis approach can be used to quantify variation and overall effects of virulence factors across host environments. As a test case, we focus on pyoverdine, a siderophore secreted by the opportunistic pathogen *Pseudomonas aeruginosa* to scavenge iron from the host environment (19). Table 1 provides an overview of the workflow of our meta-analysis, where we combined the outcomes of 76 individual virulence experiments from 23 studies (12, 20–41, see also Tables S1 and S2 in the supplemental material). Using a weighted meta-analysis approach, we were able to investigate the evidence for pyoverdine’s contribution to virulence across eight host species, including vertebrates, invertebrates and plants, five tissue infection models and various *P. aeruginosa* genotypes. We chose pyoverdine production as the model trait for our analysis because: (i) it has been extensively studied across a range of *Pseudomonas* strains (42); (ii) its virulence effects have been examined in a large number of host species; (iii) *P. aeruginosa* is one of the most troublesome opportunistic human pathogens, responsible for many multi-drug resistant nosocomial infections (43, 44); and (iv) multiple anti-virulence drugs have been proposed to target pyoverdine production and uptake (22, 23, 45). The applied question here, then, is whether targeting pyoverdine could generally and effectively curb pathogenicity.

**TABLE 1.**
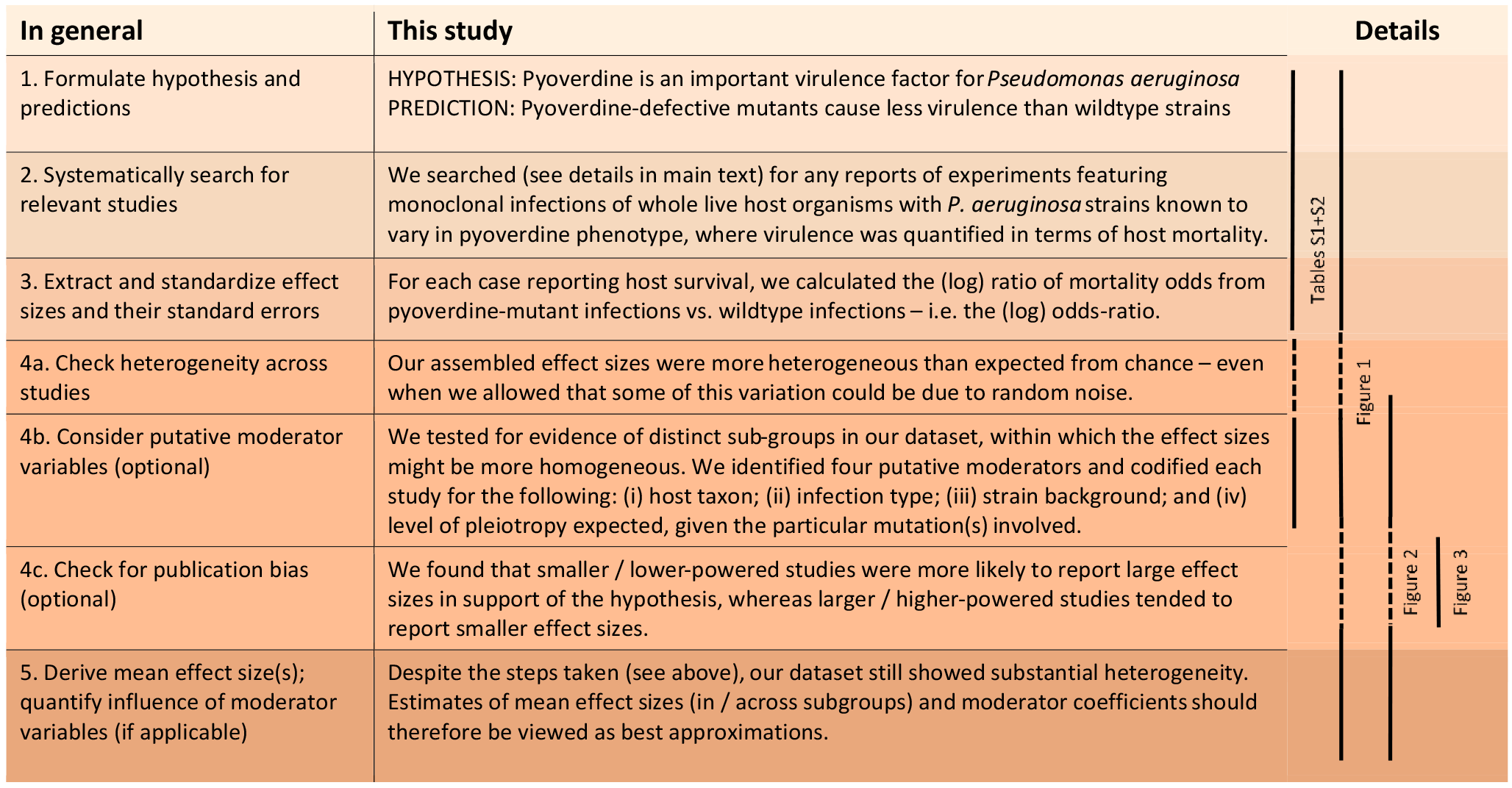
Meta-analysis workflow for this study

## RESULTS

**Literature search and study characteristics**. We searched the literature for papers featuring infections of whole live host organisms with *P. aeruginosa* strains known to vary in pyoverdine phenotype. Following a set of inclusion/exclusion rules (see materials and methods for details), we were able to include data from a total of 76 experiments from 23 original papers in our meta-analysis (Table 1; see also Fig. S1 and Tables S1 and S2 in the supplemental material). These experiments featured a range of host organisms, including mammals (mice and rabbits, n = 32), the nematode *Caenorhabditis elegans* (n = 32), insects (fruit fly, silk worm and wax worm, n = 8) and plants (wheat and alfalfa, n = 4). Experiments further differed in the way infections were established and in the organs targeted. The most common infection types were gut (n = 34), systemic (n = 16), respiratory (n = 8) and skin infections (n = 6), but we also included some other types of infections (n = 12). Each experiment compared infections with a control *P. aeruginosa* strain (which produced wildtype levels of pyoverdine) to infections with a mutant strain defective for pyoverdine production. The most common control strains used were PAO1 (n = 53) and PA14 (n = 19), which are both well-characterized clinical isolates. However, some experiments used less well-characterized wildtype strains, such as FRD1 (n = 2) and PA06049 (n = 2). Twenty-six experiments used mutant strains with clean deletions or transposon Tn5 insertions in genes encoding the pyoverdine biosynthesis pathway. In these cases, pleiotropic effects are expected to be relatively low – i.e. presumably only pyoverdine production was affected. The other 50 experiments used mutants where pleiotropic effects were likely or even certain. For example, some mutant strains carried mutations in *pvdS*, which encodes the main regulator of pyoverdine production that also regulates the production of toxins and proteases (46, 47). Others carried mutations in *pvdQ*, encoding an enzyme known to degrade quorum-sensing molecules in addition to its role in pyoverdine biosynthesis (27).

**Relationship between effect sizes and moderator variables**. We combined data from the set of experiments described above in a meta-analysis to determine the extent to which pyoverdine’s effect on virulence varied across four moderator variables: (i) host taxa, (ii) tissue types, (iii) pathogen wildtype background, and (iv) pyoverdine-mutation type. To obtain a comparable measure of virulence across experiments, we extracted in each instance the number of cases where a given infection type did or did not have a virulent outcome (i.e. dead vs. alive, or with vs. without symptoms) for both the mutant (*m*) and the wildtype (*w*) strain for each experiment (see materials and methods for details). We then took as our effect size the log-odds-ratio, i.e. ln ((*m*_virulent_ / *m*_non-virulent_) / (*w*_virulent_ / *w*_non-virulent_)) (see Table S2 in the supplemental material), a commonly-used measure especially suitable for binary response variables like survival (48).

Consistent with the theoretical prediction that host-pathogen interactions and host ecology are important modulators of virulence, we found considerable variation in the effect sizes across experiments and subgroups of all moderators (Fig. 1). Pyoverdine-deficient mutants showed substantially reduced virulence in invertebrate and mammalian hosts, whereas there was little evidence for such an effect in plants (Fig. 1A). Overall, evidence for pyoverdine being an important virulence factor was weak for taxa with a low number of experiments (i.e. for plants, and the insect models *Drosophila melanogaster* and *Galleria mellonella*). We found that pyoverdine-deficient mutants exhibited reduced virulence in all organs and tissues tested, with the exception of plants (Fig. 1B). Comparing the effect sizes across wildtype strain backgrounds, we see that pyoverdine deficiency reduced virulence in experiments featuring the well-characterized PA14 and PAO1 strains (Fig. 1C) whereas the reduction was less pronounced in experiments with less well-characterized wildtype strains. This could be due to sampling error (only a few experiments used these strains) or it may be that these strains really behave differently from PA14 and PAO1. Finally, we observed that the nature of the pyoverdine-deficiency mutation matters (Fig. 1D). Infections with strains carrying well-defined mutations known to exclusively (or at least primarily) affect pyoverdine production showed a relatively consistent reduction in virulence. Conversely, where mutants were poorly-defined, or carried mutations likely to affect other traits beyond pyoverdine, here the virulence pattern was much more variable, with both reduced and increased virulence relative to wildtype infections (Fig. 1D). We posit that at least some of the differences in observed virulence between these mutants and their wildtype counterparts was likely due to pleiotropic differences in phenotypes unrelated to pyoverdine.

**FIG. 1.**
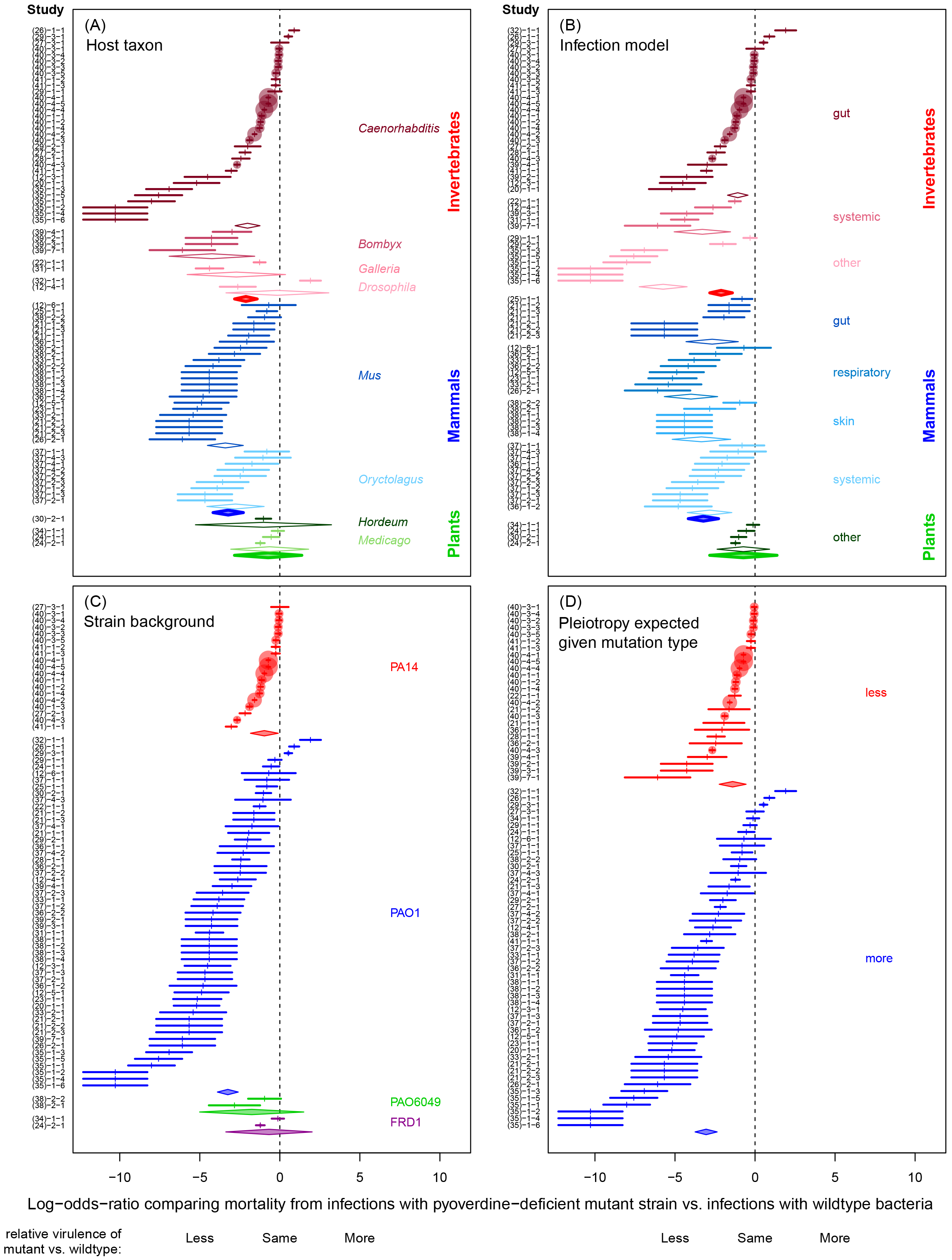
Forest plots depicting the variation in effect size across experiments on pyoverdine as a virulence factor in *P. aeruginosa*. All panels display the same effect sizes originating from the 76 experiments involved in the meta-analysis, but grouped differently according to four moderator variables, which are: (A) host taxon; (B) infection type; (C) wildtype strain background; and (D) the likelihood of pleiotropy in the pyoverdine-deficient strain. Effect sizes are given as log-odds-ratio ± 95% confidence interval. Negative and positive effect sizes indicate lower and higher virulence of the pyoverdine-deficient mutant relative to the wildtype, respectively. Diamonds represent the mean effect sizes (obtained from meta-regression analysis) for each subgroup of a specific moderator variable. IDs of the individual experiments are listed on the Y-axis (for details, see Table S1 in the supplemental material). The numbers in brackets on the Y-axis correspond to the citation number of the corresponding publication.

**Assessing the relative importance of moderator variables**. Fig. 1 highlights that we are dealing with an extremely heterogeneous dataset (a random metaanalyses of the full dataset without moderators yielded heterogeneity measures *I*^2^ = 98.1% and *H* = 7.28). Much of the variation we observe is probably due to other factors beyond those explored in Fig 1. The issue is that (a) we do not know what all these additional factors might be, and (b) the probably patchy distribution of experiments across the levels and ranges of these other factors would leave us with limited power to test for their effects. Accordingly, we decided to focus our attention on quantifying the impact of the four previously described moderators by using a more homogenous core dataset (n = 50), where rare and poorly characterized subgroups were removed. Specifically, we excluded experiments involving plants and / or undefined wildtype strains (n = 6), experiments reporting tissue damage as a measure of virulence (n = 12), and experiments where the hosts were likely not colonized by bacteria but died from exposure to bacterial toxins (n = 8). This leaves us with a core dataset comprising only those experiments where animal host models were infected with strains from well-defined PA14 or PA01 wildtype background, and survival vs. death was used as a virulence endpoint.

Using this restricted dataset, we performed a series of meta-regression models to test for significant differences between subgroups of our moderator factors, and we also estimated the share of total variance in effect sizes that is explained by each moderator variable (Fig. 2). These models revealed that infection type is the variable that explains the largest share of total variance (25.6%). For instance, in systemic infection models the pyoverdine-defective mutants showed strongly reduced virulence compared to the wild-type, whereas this difference was less pronounced in gut infections. Host taxon explained only 7.9% of the total variance in effect sizes, and there was no apparent difference in the mean effect size among invertebrate vs. mammalian host models. Finally, the wildtype strain background and the likelihood of pleiotropy in the mutant strain both explained less than 1% of the overall effect size variation, and accordingly, there were no apparent differences between subgroups (Fig. 2). Note that even with the inclusion of these moderator factors in the model, substantial heterogeneity remained in our restricted data set (*I*^2^ = 96.8%, *H* = 5.60).

**FIG. 2.**
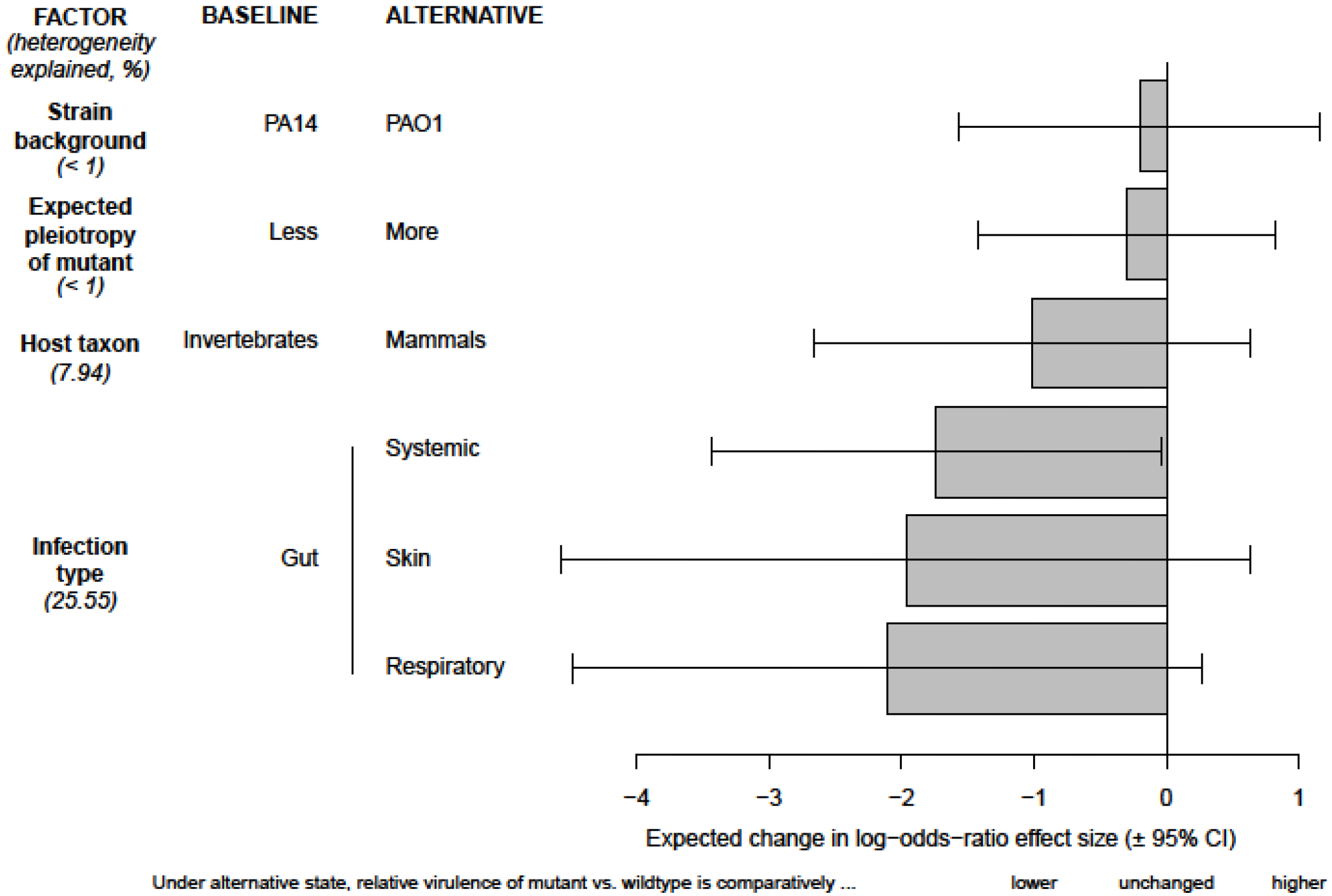
Test for differences between subgroups of moderator variables with regard to the effect sizes for pyoverdine as a virulence factor in *P. aeruginosa*. Our baseline condition for all comparisons is the following: gut infections in invertebrate hosts, using the *P. aeruginosa* wildtype strain PA14 vs a pyoverdine-deficient PA14 mutant with a low expected level of pleiotropy. The effect size for this baseline scenario is set to zero. All other scenarios had more extreme (negative) effect sizes, and are therefore scaled relative to this baseline condition. Comparisons reveal that virulence in pyoverdine-deficient strains was significantly more reduced in systemic compared to gut infections, and that most effect size variation is explained by the infection type. There were no significant effect size differences between any of the other subgroups. Bars show the difference in log odds-ratio (± 95% confidence interval) between the baseline and any of the alternate conditions. Values given in brackets indicate percentage of effect size heterogeneity explained by a specific moderator.

**Publication bias**. In any field, there is a risk that studies with negative or unanticipated results may be less likely to get published (e.g. in our case, pyoverdine-deficient mutants showing no change or increased levels of virulence) (49). Especially when negative or unanticipated results are obtained from experiments featuring low sample sizes (and thus high uncertainty), the scientists responsible may be less inclined to trust their results, and consequently opt not to publish them. This pattern could result in a publication bias, and an overestimation of the effect size. To test whether such a publication bias exists in our dataset, we plotted the effect size of each experiment against its (inverted) standard error (Fig. 3). If there is no publication bias, we would expect to see an inverted funnel, with effect sizes more or less evenly distributed around the mean effect size, irrespective of the uncertainty associated with each estimate (i.e. position on the y-axis). Instead, we observed a bias in our dataset, with many lower-certainty experiments that show strongly negative effect sizes (i.e. supporting the hypothesis that pyoverdine is important for virulence; Fig. 3) but a concomitant paucity of lower-certainty experiments that show weakly negative, zero or positive effect sizes (i.e. not supporting the hypothesis).

**FIG. 3.**
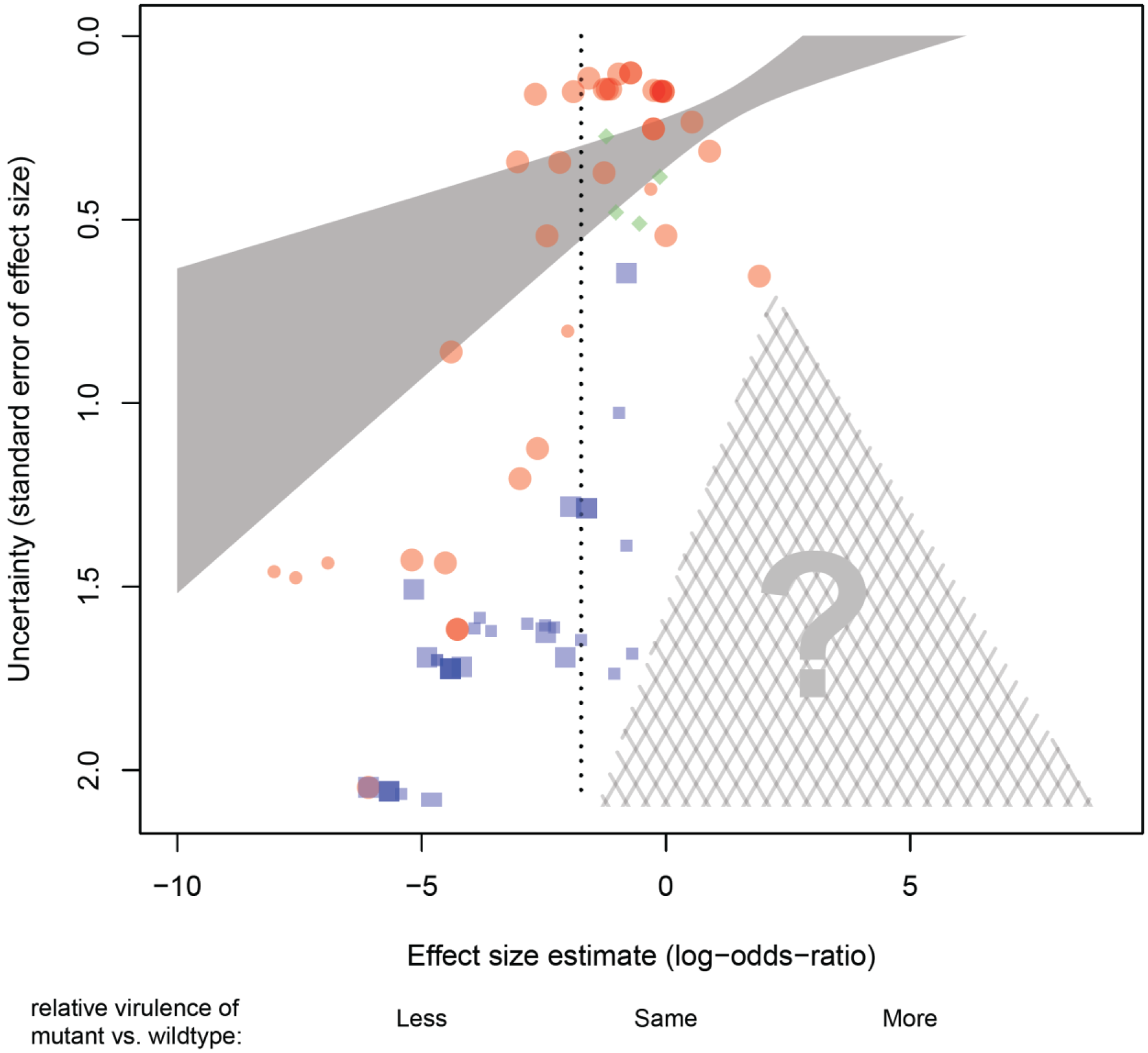
Association between effect sizes and their standard errors across 76 experiments examining the role of pyoverdine production for virulence in *P. aeruginosa*. In the absence of bias, we should see an inverted funnel-shaped cloud of points, more or less symmetrically distributed around the mean effect size (vertical dotted line). Instead, we see an over-representation of low-certainty experiments associated with strong (negative) effect sizes. This suggests a significant publication bias: experiments with low-certainty and weak or contrary effects presumably do exist, but are under-represented here (note the absence of data points in the cross-hatched triangle). Effect sizes are given as log-odds-ratio. Each symbol represents a single experiment. Symbol colours and shapes stand for different host organisms (red circles = invertebrates; blue squares = mammals; green diamonds = plants). Large symbols denote the experiments included in the core dataset. The solid shaded area represents the 95% confidence interval for the weighted linear regression using the complete dataset. Note that due to the stronger weights accorded to high certainty experiments (i.e. the points towards the top of the plot), many of the lower-weighted (higher-uncertainty) points towards the bottom of the plot lie quite far from the regression line and also outside the confidence interval.

## DISCUSSION

**What we can conclude from this meta-analysis**. Our meta-analysis reveals that pyoverdine-deficient strains of the opportunistic pathogen *P. aeruginosa* typically showed reduced virulence across a wide range of host species and bacterial genotypes. This confirms that iron limitation is a unifying characteristic of the host environment, making siderophores an important factor for pathogen establishment and growth within the host (50, 51). Conversely, we also saw that the extent to which pyoverdine deficiency reduced virulence varied considerably, and was quite modest in many instances. Pyoverdine-deficient mutant strains were typically more benign, owing to a reduced capacity for *in vivo* growth and/or a reduced capacity for inflicting damage on their host. Nonetheless, these mutants were typically still able to establish a successful infection, and, in many cases, could still kill their host (21, 22, 25, 40). These results support ecological theory predicting that the effect of a certain phenotype (i.e. producing pyoverdine in our case) should vary in response to the environment (i.e. the host and infection context). Our findings have consequences for any therapeutic approaches targeting this particular virulence factor as they reveal a possible trade-off: such treatments could have wide applicability, but their (clinical) impact would likely vary across infection contexts, and be limited to attenuating rather than curing the infection. This would mean that for *P. aeruginosa* infections, at least, therapies targeting siderophore production could be helpful but should probably still be accompanied by other therapeutic measures (52).

Our work demonstrates how meta-analyses can be used to quantitatively synthesize data from different experiments carried out at different times by different researchers using different designs. Such an analytical approach goes beyond a classical review, where patterns are typically summarized in a qualitative manner. For instance, a recent study proposed that three different virulence factors (pyocyanin, protease, swarming) of *P. aeruginosa* are host-specific in their effects (12). Here we use a meta-analytic approach to quantitatively derive estimates of the overall virulence potential of a given bacterial trait and investigate variables that affect infection outcomes. We assert that such quantitative comparisons are essential to identify those virulence factors that hold greatest promise as targets for effective broad-spectrum anti-virulence therapies.

Our finding that effect sizes vary considerably across our assembled experiments provides a different perspective compared to that which one would obtain from a cursory reading of the literature. For instance, the first study investigating pyoverdine in the context of an experimental infection model (38) reported that pyoverdine is essential for virulence. Although this experiment and its message have been widely cited (including by ourselves), it may no longer be the strongest representative of the accumulated body of research on this topic. As we see in Figure 1, the effect size it reports is associated with a high uncertainty due to a comparatively low sample size. Moreover, the observed effect cannot unambiguously be attributed to pyoverdine because an undefined UV-mutagenized mutant was used. We highlight this example not to criticise it, but rather because it serves to demonstrate why drawing inferences from (appropriately weighted) aggregations of all available evidence is preferable to focusing solely on the results of a single study.

**What we could conclude with additional data**. Our meta-analytic approach not only provides information on the overall importance of pyoverdine for *P. aeruginosa* virulence, but it also allows us to identify specific gaps in our knowledge and potential biases in the published literature. First, most experiments in our dataset employed acute infection models, even though *P. aeruginosa* is well known for its persistent, hard-to-treat chronic infections. This raises the question to what extent insights on the roles of virulence factors important in acute infections can be transferred to chronic infections. In the case of pyoverdine, we know that in chronically-infected cystic fibrosis airways, pyoverdine production is often selected against (54–56). Although the selective pressure driving this evolutionary loss is still under debate (current explanations include pyoverdine disuse, competitive strain interactions and/or a switch to alternative iron-uptake systems (56–58)), this example illustrates that the role of pyoverdine might differ in acute versus chronic infections.

Second, our comparative work shows that experiments were predominantly carried out with the well-characterized strains PAO1 and PA14. While these strains were initially isolated from clinical settings, they have subsequently undergone evolution in the laboratory environment (59–61), and might now substantially differ from the clinical strains actually causing acute infections in hospitals. Therefore, while we found no overall differences between the lab strains used in our data set, we argue that it would still be useful to carry out additional studies on a range of clinical isolates to be able to make firm conclusions on the general role of pyoverdine as a virulence factor.

Finally, our data analysis revealed that low-certainty studies showing no or small effects of pyoverdine on virulence were under-represented in our data set, which points towards a systematic publication bias. It remains to be seen whether such biases are common with regard to research on virulence factors, and whether they result in a general overestimation of the effect these factors have on host survival or tissue damage. With regard to pyoverdine, further studies are clearly needed to obtain a more accurate estimate of the true effect size.

**Guidelines for future studies**. While our study demonstrates the strength of quantitative comparative approaches, it is important to realize that extracting effect sizes is one of the biggest challenges in any meta-analysis. This challenge was particularly evident for the experiments we found, which profoundly varied in the way data was collected and reported. As a consequence, we had to exclude many studies because they used measures of virulence that were only reported by a minority of studies, or because their reporting of results was unclear (for a selected list of examples, see Table S3 in the supplemental material). To amend this issue for future studies, we would like to first highlight the problems we encountered and then provide general guidelines of how data reporting could be improved and standardized. One main problem we experienced was incomplete data reporting (i.e. mean treatment values, absolute values and/or sample size was not reported), which prevents the calculation of effect sizes and uncertainty measures. Another important issue was that different studies measured virulence using very different metrics. Some measured virulence at the tissue level (i.e. the extent of damage inflicted), while others focused on the whole host organism. Others focused on the dynamics of the bacteria themselves, taking this as a proxy for the eventual damage to the host. There were both quantitative measures (e.g. extent of damage), and qualitative measures (e.g. assignments to arbitrary categories of virulence). Survival data was sometimes presented as a timecourse, sometimes as an endpoint; sometimes as raw counts, sometimes as proportions. In most cases, the time scales over which survival was assessed were fairly arbitrary. Compiling such diverse measures of virulence is not simply time consuming, but it also generates extra sources of heterogeneity in the dataset, which might interfere with the basic assumptions of meta-analytical models (62, 63).

How can these problems be prevented in future studies? We propose the following. (a) Whenever possible, time-to-event data (e.g. death, organ failure, etc.) should be recorded in a form that preserves both the outcome and the times to event per subject. (b) The number of replicates used (hosts) and a measure of variance among replicates must be provided to be able to calculate a confidence estimate for the experiment. (c) If data are scaled in some way (e.g. relative to a reference strain), the absolute values should still be reported, because these are crucial for the calculation of effect sizes. Finally, (d) studies leading to unexpected or negative results (e.g. no difference in virulence between a wildtype and a mutant) should still be published, as they are needed to estimate a true and unbiased effect size. In summary, all findings, irrespective of their magnitude or polarity, should be presented “as raw as possible” (e.g. in supplementary files or deposited in online data archives). This will make comparisons across studies much easier and will provide a useful resource for future meta-analytic studies.

**Conclusions**. Currently, bacterial traits are subject to a binary categorisation whereby some are labelled as virulence factors while others are not. We demonstrate that traits’ effects on virulence are anything but binary. Rather, they strongly depend on the infection context. Our study affirms meta-analysis as a powerful tool to quantitatively estimate the overall effect of a specific virulence factor and to compare its general importance in infections across different bacterial strains, hosts, and host organs. Such quantitative comparisons provide us with a more complete picture on the relative importance of specific virulence factors. Such knowledge is especially valuable for opportunistic pathogens, which have a wide range of virulence factors at their disposal, and infect a broad range of host organisms (13–15). Meta-analytical comparisons could thus inform us on which traits would be best suited as targets for anti-virulence therapies. Ideal traits would be those with high effect sizes and general importance across pathogen and host organisms.

## MATERIALS AND METHODS

**Literature search**. We conducted an extensive literature search, using a combination of two online databases: Web of Science and Google Scholar. The following terms were used to search abstracts and full texts: “aeruginosa” in combination with “pyoverdin” or “pyoverdine” and in combination with “virulence” or “infection” or “pathogen” or “disease” or “mortality” or “lethality”. This search was first performed on May 19^th^ 2014 and it was repeated periodically until Sep 21^st^ 2015 in order to include more recent publications. In addition, the reference lists of all shortlisted studies were scanned for relevant publications. We further contacted the corresponding authors of several publications to ask for unpublished datasets.

**Inclusion criteria**. The database search yielded a total of 442 studies, and we identified 10 additional records through other sources. These 452 studies were then scanned for relevant content according to the following set of inclusion criteria. Studies were considered potentially eligible for inclusion if they contained original research, were written in English and provided data that compared the virulence of a wildtype pyoverdine-producing *P. aeruginosa* strain with that of a mutant strain demonstrating impaired pyoverdine production. We defined virulence as a decrease in host fitness, measured as an increase in mortality or tissue damage when infected with bacteria. We defined “wildtype strains” as strains that were originally clinical isolates, have been widely used in laboratories as virulent reference strains and have not been genetically modified. Strains with impaired pyoverdine production included strains that were completely deficient in pyoverdine production and strains that were only partially deficient, i.e. that produced less than the wildtype strain under identical experimental conditions. We considered both genetically engineered knock-out strains and clinical isolates with reduced pyoverdine production. There were 31 original publications containing 115 experiments that satisfied these criteria and were thus considered appropriate for in-depth examination.

We screened these 115 experiments using a second set of rules to identify those experiments that contain comparable quantitative data, which is essential for a metaanalysis. Inclusion criteria were: (i) virulence was measured directly (and not inferred indirectly via genetic analysis); (ii) virulence was measured *in vivo* (and not *in vitro* via virulence factor production); (iii) virulence was measured quantitatively as direct damage to the host caused by bacterial infections, and not by indirect or qualitative measures such as bacterial growth performance in the host, threshold infective dose required to kill a host, the damage associated with virulence factor administration, or resistance to macrophage-like predation (53, 64, 65); and (iv) absolute virulence data were presented (and not only data scaled relative to the wildtype without information on the absolute risk of mortality, since effect sizes cannot be calculated from such data). This second set of rules was fulfilled by 23 original publications containing 76 individual experiments (see Tables S1 and S2 in the supplemental material). For an overview of the whole selection process, see Fig. S1 in the supplemental material.

**Data extraction and effect size calculations**. From all of these 76 experiments, we extracted information on: (i) the host organism; (ii) the type of infection; (iii) the observation period of infected hosts; (iv) the identity of the control (wildtype) strain; (v) the identity of the pyoverdine-deficient strain; (vi) the mutated gene in the pyoverdine-defective strain; (vii) the mutation type (e.g. insertion/deletion); (viii) the sample size used for the wildtype and mutant experiments; and (ix) the relevant virulence measure (host survival or tissue damage) for wildtype and mutant strains (see Table S1 in the supplemental material). This information was used to categorize the experiments and identify potentially important moderator variables (see below).

Next, we extracted quantitative data from these experiments so we could calculate effect sizes for the virulence associated with pyoverdine production. For mortality assays, we extracted raw counts of how many individuals died and how many survived following infection with a dose of *P. aeruginosa* wildtype or, alternatively, a mutant strain known to be deficient for pyoverdine production. For experiments on tissue damage, we extracted information on the number of individuals with and without the symptoms related to tissue damage (e.g. a lesion in an organ). In cases with zero counts (i.e. either all or none of the individuals in a particular treatment group died or experienced tissue damage), we converted counts to 0.5 to avoid having zero denominators in the subsequent calculations of the (log-odds ratio) effect sizes (66). In cases where data from multiple time-points or survival curves were available, we concentrated on the time point with the largest difference between the wildtype and the mutant infection.

Using this count data, we calculated the effect size for each experiment as the log-odds-ratio = ln ((*m*_virulent_ / *m*_non-virulent_) / (*w*_virulent_ / *w*_non-virulent_)), where *m*_virulent_ and *W*_virulent_ are the number of individuals that died or experienced tissue damage when infected by the mutant and the wildtype strain, respectively, and *m*_non-virulent_ and Wnon-virulent are the number of individuals that survived or remained unharmed by the infection. Information on the sample size was used to calculate the 95% confidence interval for each effect size and for weighting effect sizes relative to one another (see details below). Where experiments reported a range of sample sizes, we used the arithmetic mean. Some studies reported only a minimum sample size. In those cases, we used this number. For experiments using *C. elegans,* infections were often carried out on replicate petri dishes in a large number of individuals. In these cases, we used the total number of individual worms used in each treatment group as sample size, and not the number of replica plates.

For some experiments conducted in mammals, data on *in vivo* growth of a wildtype and a pyoverdine deficient strain were available in addition to virulence measures (see Table S4). We also calculated effect sizes (standardized mean differences) for this set of studies (Fig. S4). This limited dataset shows a similar pattern to the main dataset shown in Fig. 1, but was not included in the main analysis because we were primarily interested in quantifying virulence effects (i.e. host damage and/or mortality) and not pathogen growth.

**Moderator variables**. We considered four moderator variables (host taxon, infection type, wildtype strain background, likelihood of pleiotropy associated with pyoverdine deficiency) that could potentially explain variation in virulence. In cases where information was missing for a specific moderator variable, we contacted the authors to obtain additional information. For each moderator, we defined the following relevant subgroups.

*Host organism* – We first split experiments into broad taxonomic units (mammals, invertebrates, plants), and then classified hosts by genus.

*Infection type* – We classified experiments according to the organ or body region targeted by the infection. Major categories include infections of the host organisms’ respiratory tract, digestive system, skin (including burn wounds), and infections that generated a non-localized infection of the body cavity (systemic infection). Experiments that did not fit in any of these categories, such as infections of whole seedlings, were classified as ‘other infection types’.

*Wildtype strain background* – Four different *P. aeruginosa* wildtypes (PAO1, PA14, FRD1 and PAO6049) were used for infection experiments. Although it is well established that even standard strains such as PAO1 can substantially differ between labs, there was not enough information available to take such strain-level variation into account.

*Likelihood of pleiotropy* – The focal phenotype investigated in this meta-analysis is the production of pyoverdine, the main siderophore of *P. aeruginosa*. Mutants exhibiting reduced or no pyoverdine production can be generated either by deleting a specific pyoverdine-synthesis gene, or through untargeted mutagenesis (e.g. UV light)(67). The latter mutants are likely to have mutations in other genes unrelated to pyoverdine synthesis. These mutations are typically unknown but could also affect virulence. In principle, even single gene deletions can have pleiotropic effects on the phenotype, via disruption of interactions with other genes. Depending on the locus in question, certain genetic modifications are more likely to induce pleiotropy than others. To account for these complications, we inferred on a case-by-case basis whether the mutation used was likely to only induce a change in (or loss of) pyoverdine production (i.e. pleiotropy less likely) or was likely to induce a change in other phenotypes as well (i.e. pleiotropy more likely). In the biosynthesis of pyoverdine, multiple enzymes are involved in non-ribosomal peptide synthesis (19). Two gene clusters, the *pvc* operon and the *pvd* locus, encode proteins involved in the synthesis of the chromophore and peptide moieties, respectively (19, 68). In most of these genes, a mutation or deletion leads to a complete loss of pyoverdine production, and most likely does not affect any other trait. Accordingly, we assigned mutants carrying mutations in these genes to the category “pleiotropy less likely”. An exception is *pvdQ*, a gene coding for a periplasmic hydrolase, which is required for pyoverdine production, but is also involved in the degradation of N-acyl-homoserine lactone quorum-sensing molecules (27). Strains with deletions in this gene were therefore assigned to the category “pleiotropy more likely”. Other strains falling into this category included: (i) mutants where the key regulator of pyoverdine synthesis, PvdS, was deleted, leading to deficiencies in toxin and protease production, in addition to a complete loss of pyoverdine production (69); (ii) strains that carry a deletion in a central metabolic gene and only coincidentally show no (or strongly reduced) pyoverdine production; (iii) double mutants that carry deletions in both the pyoverdine and the pyochelin synthesis pathway (pyochelin is the secondary siderophore of *P. aeruginosa*) (70); and (iv) pyoverdine mutants created via non-targeted (e.g. UV) mutagenesis.

**Core dataset**. To quantify the impact of the moderator variables, we removed experiments belonging to rare or poorly characterized subgroups to generate a more homogenous core dataset. We excluded experiments involving plants and / or undefined wildtype strains (n = 6), experiments reporting tissue damage as a measure of virulence (n = 12), and experiments where the hosts were likely not colonized by bacteria but died from exposure to bacterial toxins (n = 8). This resulted in a core dataset comprising 50 experiments that was used for subsequent analyses on the influence of moderator variables.

**Statistical analysis**. Analyses were performed in R version 3.2.3 (71), using functions from packages ‘meta’ (72) and ‘metafor’ (73). We used the ‘metabin’ function to transform the count data into the (log-) odds ratio described above. We then weighted these values by the inverse of their respective squared standard errors, and pooled them to obtain a single distribution of effect sizes. We reasoned that the variability of the effect sizes in our dataset probably reflects more than simple sampling error around a single true mean. Rather, we assume that our effect sizes represent a random sample from a larger distribution comprising all possible true effect size estimates. As such, we inferred that a random effects meta-analysis would be more appropriate for our dataset than a fixed effects model (for further discussion, see (62, 63). In a random effects meta-analysis, we partition the total heterogeneity observed in our dataset (described by the statistic Q) into two constituent parts – within-experiment variation (ɛ) and between-experiment variation (ζ). The latter component, scaled appropriately to account for the weightings intrinsic to metaanalysis, is quantified as the *T^2^* statistic. There are several different algorithms one can use to effect this partitioning of variance. We chose a restricted maximum likelihood (REML) approach. The use of a random model, rather than a simpler fixed model, affects the weights accorded to each constituent effect size, which in turn changes our estimates for pooled means and their associated errors. We further slightly broadened confidence intervals and weakened test statistics using Knapp and Hartung’s algorithm (74) – a widely-used and conservative adjustment designed to account for the inherent uncertainty associated with the partitioning of heterogeneity we perform in the course of fitting a random effects model.

We assessed the degree of residual heterogeneity in our dataset using statistics *I^2^* and *H. I^2^* estimates the approximate proportion of total variability across experiments that is attributable to unexplained heterogeneity, as opposed to simple sampling error (chance). *H* reports ‘excess’ heterogeneity as a fold difference compared to the baseline amount of variability we would have expected if the sample were homogenous (75).

Both metrics described above indicated considerable residual heterogeneity in our dataset, so we suspected that, beyond the random-and sampling error, some measurable characteristics of the experiments in our dataset could be contributing, in predictable ways, to the observed heterogeneity of our assembled effect sizes. We investigated four potential moderators, namely the host taxon, the type of infection, the wildtype background of the infecting strains, and the type of mutant involved (i.e. whether more or less pleiotropy was expected). In a first approach, we split the dataset into subgroups representing different levels of these moderators, and estimated pooled means within these different subgroups. For this, we again used random-effect meta-analysis (as above), but we set the level of between-experiment heterogeneity (*T^2^*) to be common across all subgroups.

In a second approach, we fitted a series of meta-regression models that extended our basic model to additionally consider the contributions of multiple moderator factors. Our models were able to estimate moderators’ additive effects only, because the distribution of data across different combinations of factor levels was too patchy to permit a proper investigation of moderators’ interactive effects. Moderators’ alterations of the expected (i.e. baseline) effect size could be quantified as coefficients, which could, when standardized as t-statistics, be tested for significant differences from zero. In addition, we could test whether, collectively, the inclusion of moderators in our meta-analysis model significantly reduced the residual heterogeneity relative to a situation with no moderators.

To estimate what share of the residual heterogeneity in our dataset could be individually attributable to each of the respective moderators, we performed a series of likelihood ratio tests comparing, in each case, a full model including all four moderators, against a reduced model that excluded one of the moderators. Variance component estimation in these models used maximum likelihood instead of REML because nested REML models cannot be compared in this way. From each pairwise comparison, we obtained a pseudo-R^2^ value, which reflects the difference in *T^2^* (between-experiment heterogeneity) between the two models, scaled by the *T^2^* of the simpler model.

To test for putative publication bias in our dataset, we compared effect sizes against their respective standard errors, the idea being that if there is no bias, there should be no link between the magnitude of the result from a given experiment, and the ‘noisiness’ or uncertainty of that particular result. If there is bias, we could find an overrepresentation of noisier experiments reporting higher magnitude results. Using the ’metabias’ function of the R package ‘meta’, we performed both (weighted) linear regressions and rank correlations to test for this pattern (76, 77).

## ACKNOWLEDGEMENTS

We thank Natasha Kirienko and Kendra Rumbaugh for sharing raw and unpublished data and Richard Allen, Steve Diggle, Sinead English, Steven Higgins and Shinichi Nakagawa for comments on the manuscript.

## AUTHOR CONTRIBUTIONS

E.G., F. H., R.K., and A.R.G. conceived the study; E.G. conducted the literature search and compiled the data set; A.R.G. conducted statistical analysis; E.G., F.H., R.K., and A.R.G. interpreted the data and wrote the paper.

---

**FIG. S1.**
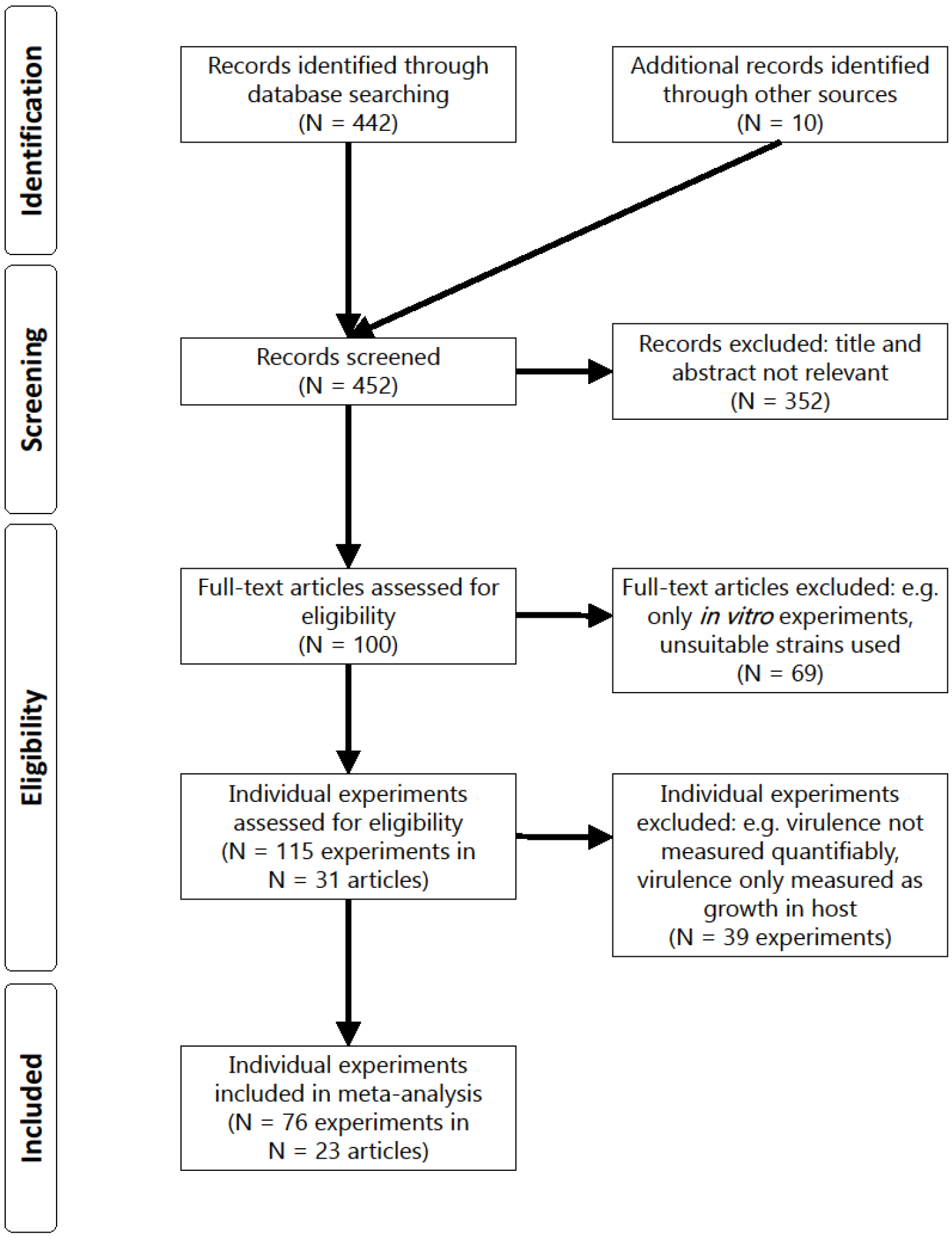
Flow diagram (PRISMA format) of the screening and selection process for studies investigating the association between pyoverdine production and virulence in *P. aeruginosa*.

**FIG. S2.**
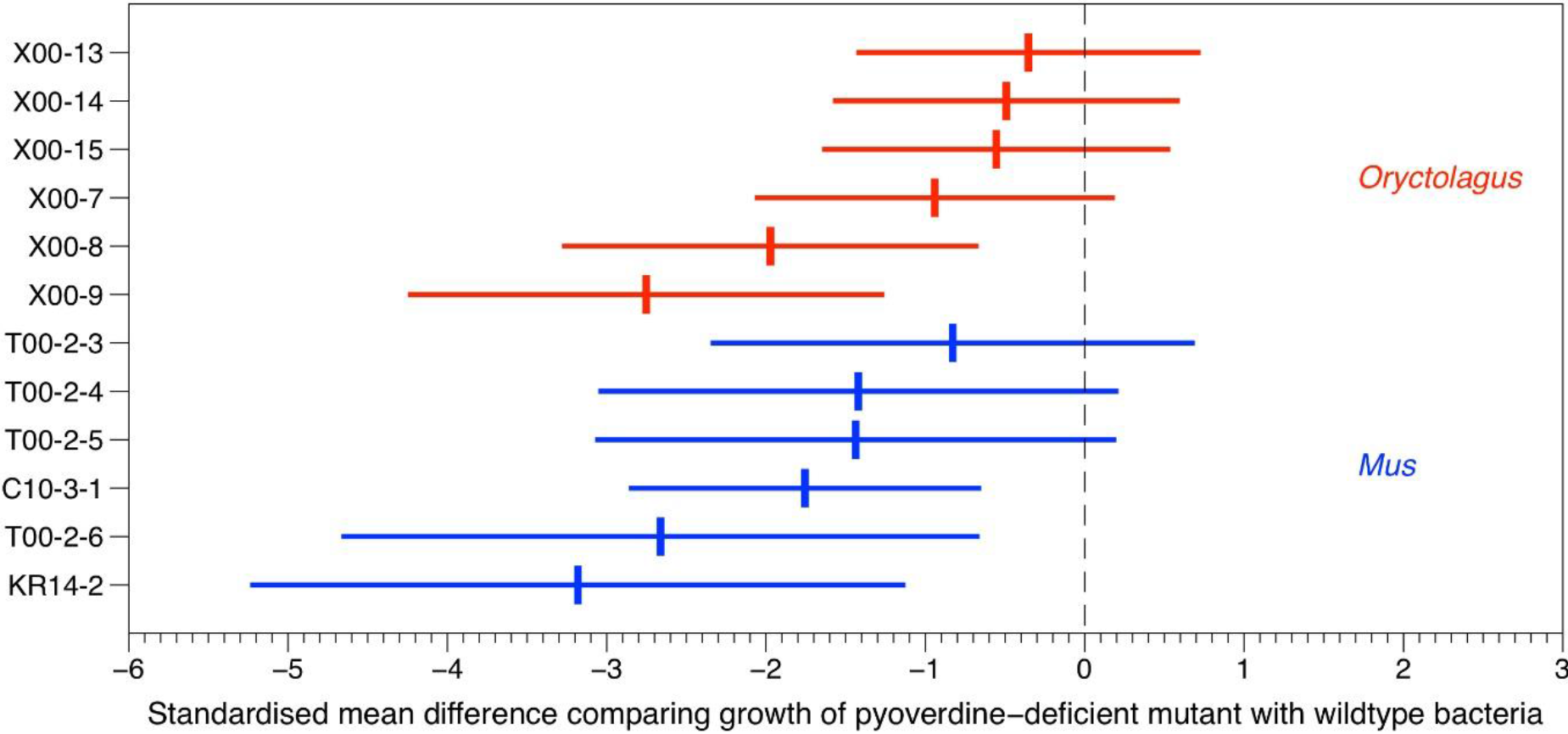
Forest plot depicting the variation in effect size across experiments on the effect of pyoverdine on the growth of *P. aeruginosa* in mammalian hosts. Effect sizes are given as standardized mean difference ± 95% confidence interval and are grouped by host genus. Negative and positive effect sizes indicate lower and higher *in vivo* growth of the pyoverdine-deficient mutant relative to the wildtype, respectively. IDs of the individual experiments are listed on the Y-axis (for details, see Table S4 in the supplemental material).

## SUPPLEMENTARY FILE LEGEND

**TABLES S1-S4** Full dataset collected for meta-analysis, including calculated effect sizes and a list of excluded experiments.

